# Identification of chromatin loops from Hi-C interaction matrices by CTCF-CTCF topology classification

**DOI:** 10.1101/2020.07.21.214585

**Authors:** Silvia Galan, François Serra, Marc A. Marti-Renom

## Abstract

Genome-wide profiling of long-range interactions has revealed that the CCCTC-Binding factor (CTCF) often anchors chromatin loops and is enriched at boundaries of the so-called Topologically Associating Domains, which suggests that CTCF is essential in the 3D organization of chromatin. However, the systematic topological classification of pairwise CTCF-CTCF interactions has not been yet explored.

Here, we developed a computational pipeline able to classify all CTCF-CTCF pairs according to their chromatin interactions from Hi-C experiments. The interaction profiles of all CTCF-CTCF pairs were further structurally clustered using self-organizing feature maps and their functionality characterized by their epigenetic states. The resulting clusters were then input to a convolutional neural network aiming at the *de novo* detecting chromatin loops from Hi-C interaction matrices.

Our new method, called LOOPbit, is able to automatically detect significant interactions with a higher proportion of enhancer-promoter loops compared to any other callers. Our highly specific loop caller adds a new layer of detail to the link between chromatin structure and function.

## INTRODUCTION

The human genome consists of 2 meters of DNA and its folding in the cell nucleus is not random (1). During the last decade, the three-dimensional (3D) organization of the genome has been associated with the regulation of multiple nuclear functions like DNA replication, repair, rearrangement and recombination, RNA processing, and transcription (1–4). From chromosome territories (5) to chromatin compartments (6), topologically associating domains (TADs) (7, 8), and finally loops (9), the hierarchical organization of the genome has been gaining in details with the outcome and further improvement of the high-throughput Chromosome Conformation Capture (3C) technologies (6). This detailed characterization of the chromatin structure also helped us to characterize the roles of specific proteins, with perhaps being the CCCTC-binding factor (CTCF) protein its most striking example.

CTCF is a transcription factor formed by 11 DNA-binding Zinc finger (ZF) it is found enriched in TAD borders and seems essential in mammalian enhancer-promoter loops (10). Its functionality is dependent on the location and the relative orientation of its binding sites. Interacting CTCF pairs tend to be organized in a convergent orientation, with the binding motifs facing each other (9). The distribution and organization of CTCF leading to TAD formation has been explained by the loop extrusion model that involves CTCF but also other protein complexes like cohesin (11, 12). Briefly, the cohesin complex, which forms a ring-shaped structure upon loading onto chromatin, extrudes chromatin resulting in a growing loop of DNA. The extrusion is blocked on both sides when cohesin encounters two CTCF in a convergent orientation. Various experiments have described the relevance of the binding motif orientation for loop formation by inverting or deleting CTCF binding sites using CRISPR/Cas9 experiments (12, 13). Moreover, a number of studies revealed the effect of CTCF depletion, resulting in a fainting of TADs and loop structures, while maintaining compartment organization. It suggests that CTCF and cohesin play a role in TAD and loop formation, but not necessarily in genome compartmentalization (14, 15).

Recently, several studies have applied Aggregate Peak Analysis (APA) (9), or pile-up methods (16) to analyze Hi-C datasets using mean signals from selected regions of interest (17–22), such as CTCF loops. At the level of single chromatin loops, their biological relevance in gene regulation, has led to the development of several loop callers. However, most of the methods present low reproducibility between biological replicates and a low correlation with different biological markers (23). Here, we propose to use self-organizing feature maps (SOFM) (24), an artificial neural network, to classify the signal from pairs of CTCF without any prior information about their topological organization. This approach allows us to obtain sub-populations of CTCF-CTCF interacting structures and to identify their epigenetic signature in an unsupervised manner. As a result, we next trained a convolutional neural network (CNN), and used the generated model in our tool, called LOOPbit, to identify chromatin loops in a robust, fast and genome-wide manner.

## MATERIALS AND METHODS

### CTCF ChIP-Seq and Hi-C data

CTCF ChIP-Seq experiments of Human B-lymphocyte cell line (GM12878) were downloaded from the ENCODE database (https://www.encodeproject.org/; dataset ids: ENCSR000DRZ, ENCSR000AKB and ENCSR000DZN) (25). After processing the peaks following the ENCODE pipelines available at https://github.com/ENCODE-DCC, only common peaks from all three experiments were kept resulting in a total of 52,844 CTCF peaks. Next, the orientation of the CTCF binding motifs was assessed by means of the MEME and FIMO motif-based sequence analysis tools (26, 27). Only peaks with a statistically significant CTCF motif were kept (p-value < 0.05), which resulted in a total of 41,816 *ChIP-Seq+motif* peaks. This filter discarded ~21% of the original detected CTCF ChIP-Seq peaks.

*In situ* Hi-C datasets for GM12878 cell line (9) were downloaded from the GEO database (**Supplementary Table 1**) and its replicates merged and parsed using the TADbit as previously described (28). Two resolution matrices (that is, at 100 kb and at 5 kb) were obtained and normalized using OneD (29) with default parameters. The 100 kb Hi-C matrices were next used to calculate chromosome compartmentalization (6, 30) using TADbit (28). The 5 kb resolution matrices were further parsed to subtract 50 kb squared submatrices centered in each axis of the matrices to any two of the 15,597 isolated CTCF *ChIP-Seq+motif* peaks (that is, ~38% of all selected peaks above). Isolated, or non-overlapping, peaks were defined as those *ChIP-Seq+motif* peaks with no other peak within a 50 kb window span from the center of the peak. This additional filter ensured that the observed signal in a Hi-C submatrix was due to a particular pair of CTCF-CTCF peaks and not multiple pairs. Finally, we subtracted a total of 130,655 submatrices between any pair of isolated CTCF-CTCF peaks spanning any distance between 45 kb and 1.5 Mb, which ensured to select most pairs within the size of a typical TAD in the human genome (~900 kb).

### Submatrix analysis, deconvolution, classification and clustering

All 130,655 submatrices between any pair of isolated CTCF-CTCF peaks were next analyzed with our Python based package called Meta-Waffle and available in GitHub (https://github.com/3DGenomes/metawaffle). Meta-Waffle takes as input a list of pairs of genomic coordinates and extracts submatrices of interactions for each pair. Next, Meta-Waffle will analyze, deconvolve and classify the submatrices into micro-clusters, or neurons, using a Self-Organizing Feature Map (SOFM) approach (24) available in http://neupy.com. Briefly, and for this application, Meta-Waffle extracted 11 by 11 submatrices (at 5 kb resolution) from an input full Hi-C interaction matrix. The submatrices values were next re-scaled between 0 and 1 using a sigmoid function. Third, the re-scaled submatrices were input to the SOFM based on a series of empirically optimized parameters including the SOFM grid size (each cell in this grid represented a SOFM neuron, we tried: 10×10, 20×20, 30×30, 40×40, 45×45, and 50×50), learning radius (*i.e.,* 1,2, and 5), standard deviation (*i.e.,* 0.5, 0.1, and 0.01), steps (*i.e.,* 1, 0.5, 0.1, and 0.01), and Epochs (*i.e.,* 30, 90, 130, and 200). Combination of the tested parameters resulted in a total of 1,152 generated SOFMs. Assessing which set of parameters results in the best submatrices classification is not trivial as there is no gold standard for the classification of CTCF-CTCF interaction submatrices. Therefore, we devised three evaluation metrics to select SOFM optimal parameters. First, a percentage of classified submatrices in order to minimize the number of singletons SOFM neurons (with only one CTCF-CTCF submatrix). Second, the variability between neurons (the more distinct the better). And third, a compartment segregation score to maximize the segregation of the two main genome compartments within the SOFM grid. Compartments were measured using the first eigenvector of a transformation of a chromosome Hi-C interaction matrix (6). Each value in this eigenvector represented the belonging to a compartment type of a given genomic bin. We used positive values for A compartments and negative values for B compartments. We defined the compartment segregation score as the absolute of the average eigenvector value of all genomic regions in a given neuron. The SOFM optimal parameters that maximized all three devised scores were: Grid size: 30×30; Learning radius: 5; Standard deviation: 0.5; Step: 0.01; and Epochs: 30. Finally, the optimal SOFM neurons were used to generate a hierarchical clustering by means of the Uniform Manifold Approximation and Projection (UMAP) (31), which can be used as an effective pre-processing step to enhance the performance of density-based clustering. The UMAP was computed using a local neighboring size of 6 for the manifold approximation, an effective minimum distance between embedded points of 0.3 and a number of epochs of 100 to increase its accuracy. The final 10 clusters of Hi-C CTCF-CTCF neurons were obtained with HDBSCAN (32) with default parameters, considering a minimum of 6 for the cluster size and a minimum of 6 neighbors for a point to be considered a core point.

### Chromatin states integration

The chromatin 15-states model for the GM12878 cell line was downloaded from the Roadmap Epigenomics Consortium (33). Following a similar previously published protocol (23), all 15 states were merged into 4 major classes: promoter (Active TSS, Flanking Active TSS, Bivalent/poised TSS Flanking bivalent TSS), enhancer (Enhancers, Genic enhancers, Bivalent enhancers), repressed polycomb (Repressed Polycomb, Weak repressed Polycomb) and heterochromatin (Heterochromatin, Quiescent/Low). Next, the genome was segmented into 5 kb bins, and each bin was classified based on the overlap (> 50 bp) with any of the four major chromatin states. A bin could be assigned to one or more categories. Next, interactions between bins were classified as “Not expected” (interacting promoter/enhancer bin with heterochromatin bin), “Promoter-Enhancer” (interacting promoter bin with an enhancer bin), and “Heterochromatin-Heterochromatin”, (interacting heterochromatin bins).

### The LOOPbit CNN

LOOPbit is a trained Convolutional Neural Network (CNN) to predict the localization of loops from Hi-C interaction matrices. LOOPbit was trained using the TensorFlow platform (https://www.tensorflow.org) with a common pattern: input matrix flattening – dense layer (ReLu) – dropout – dense layer (Softmax). As an input, the CNN takes tensors of shape (9, 9) that will be flattened in the first layer. The dense layers were used to perform the classification of input Hi-C submatrices into two different classes: loop and no-loop. The dropout layer was needed to avoid overfitting of the model. The LOOPbit CNN was trained by a 20% leave-out of the data used as test and 80% of the data for training. The training dataset obtained by sub-sampling a total of 6,000 randomly selected CTCF-CTCF submatrices from the SOFM neurons with clear signal of looping (loops) and another 6,000 randomly selected CTCF-CTCF submatrices from the SOFM neurons with no signal of looping (no-loop). In particular, loop submatrices were extracted from the HDBSCAN clusters 1, 2, 3, 4 and 5, and no-loop submatrices from HDBSCAN clusters 7, 8, 9 and 10 (**Results**). The trained model used a 0.2 drop of the data and 1,024 neurons resulting in a classification accuracy of 85.9% and 89.6% for the loop and no-loop class, respectively.

### LOOPbit benchmark

The trained model was then used to predict loop localization in 36 previously published Hi-C datasets (**Supplementary Table 2**). To avoid biases derived from the data processing pipelines, all 36 datasets were processed using the same protocol as the training datasets from GM12878 cell line (see above). Then, the Hi-C experiments were analyzed with LOOPbit, scanning chromosome-wise using a window of 9×9 bins and a step of 1 bin, to predict loops at two different resolutions (5 kb and 40 kb). The predicted localization of loops was next assessed using several metrics. First, the Jaccard Index was computed to assess reproducibility between replicates of the same experiment. Two predicted loops were considered to be identical when they shared exactly the same anchoring bins in both replicates. Second, to characterize the possible biological relevance of the predictions, the enrichment of diverse chromatin marks at loop anchors was calculated. For the benchmarking, the 15-states chromatin models for GM12878, IMR90 and h1-ESC cell lines were downloaded from the Roadmap Epigenomics Consortium (33) and analyzed as described above. For the fly late embryos dataset, the 16-chromatin states model was downloaded from modENCODE (34). As previously, the states were also merged into 4 major classes: promoter (Promoter), enhancer (Enhancer 1, Enhancer 2), repressed polycomb (PC repressed 1, PC repressed 2), and heterochromatin/Low (Heterochromatin1, Heterochromatin 2, Low signal 1, Low signal 2, Low signal 3). Similarly, the enrichment analysis of the chromatin states at the loop anchors was done as described above. Additionally, the assessment presence of CTCF sites and their orientation in the base of the predicted loops was performed only for *cis* interactions identified in the Hi-C maps at 5 kb resolution. Briefly, CTCF ChIP-seq experiments were downloaded (**Supplementary Table 3**) and peaks processed using HOMER motif analysis with default parameters. The *cis* interactions conserved in at least 2 replicates within each dataset (with exception of Jin H1-hESC with one replicate) were intersected with the list of motifs of the CTCF peaks. Then, an interaction was considered convergent if the interacting bin closer to the p-terminus of the chromosome contained one CTCF motif on the forward strand (+ orientation), and the interacting bin closer to the q-terminus of the chromosome contained one CTCF motif on the reverse strand (– orientation) (23).

## RESULTS

### Classifying CTCF-CTCF interactions using Self-Organizing Feature Maps (SOFMs) and Uniform Manifold Approximation and Projection (UMAP)

SOFMs are dependent on several parameters, such as the grid size (number of neurons or clusters), the learning radius, the step size, the standard deviation and the number of iterations or epochs (24). To unveil the best combination of SOFM parameters in the context of CTCF-CTCF submatrices classification, we first extracted the Hi-C submatrices of all possible *cis* pairs of CTCF peaks linearly separated by between 45 kb and 1.5 Mb (**Figure 1a**). These submatrices, here referred to as "CTCF-CTCF submatrices", spanned over 45 kb (20 kb of each side of the 5 kb bin with a given CTCF peak). Next, all extracted CTCF submatrices were input to a SOFM that resulted in a total of 1,152 SOFMs, each with different combinations of parameters (**Material and Methods**). To select the optimal set of parameters, we assessed three measures: (i) the percentage of classified submatrices, considering singletons (SOFM neurons with only one submatrix) as non-classified; (ii) variability between neurons; and (iii) average compartment-type segregation across neurons. These three quality measures aimed at identifying the “best” classification that maximized the number of CTCF-CTCF submatrices classified, the separation between neurons (i.e. increased variability inter-neuron), and the homogeneous compartment type within each neuron. Of all varied parameters, the SOFM grid size and the SOFM step size were the most sensitive to the final classification (**Figure 1b** and **Supplementary Fig. 1**). The three measures mentioned above were optimal for grid size of 30, learning radius of 5, standard deviation of 0.5, step of 0.01 and number of epochs of 30. Such optimal SOFM map was represented as a grid map where each cell (neuron) is composed by a set of similar CTCF-CTCF submatrices represented by its medoid submatrix (**Figure 1c**). The SOFM map clearly reflects a variety of CTCF-CTCF submatrices from loop forming pairs (lower-left corner) to non-interacting pairs (upper-right corner). Next, the resulting 900 SOFM neurons were further classified by computing the Euclidean distance between the medoid submatrices of each pair of SOFM neurons and projecting it into a two-dimensional UMAP, a non-linear manifold dimension reduction (31). We finally applied, on the UMAP coordinates of each SOFM neuron, a density-based clustering algorithm (HDBSCAN) (32). This methodology resulted in 10 clusters of SOFM neurons, each representing unique CTCF-CTCF pairing patterns. Clusters go from the most structured with the canonical cross pattern (cluster 1 including 11 SOFM neurons with a total of 2,685 CTCF-CTCF submatrices) to a completely flat pattern with no interaction (cluster 10 with 7 SOFM neurons and 1,565 CTCF-CTCF submatrices) (**Figure 1d**). Our results suggest that CTCF-CTCF pairs can adopt a variety of well-defined topological signatures observed in a Hi-C matrix. Next, we set ourselves to functionally characterize each of the detected clusters of CTCF-CTCF pairs.

**Figure 1.**
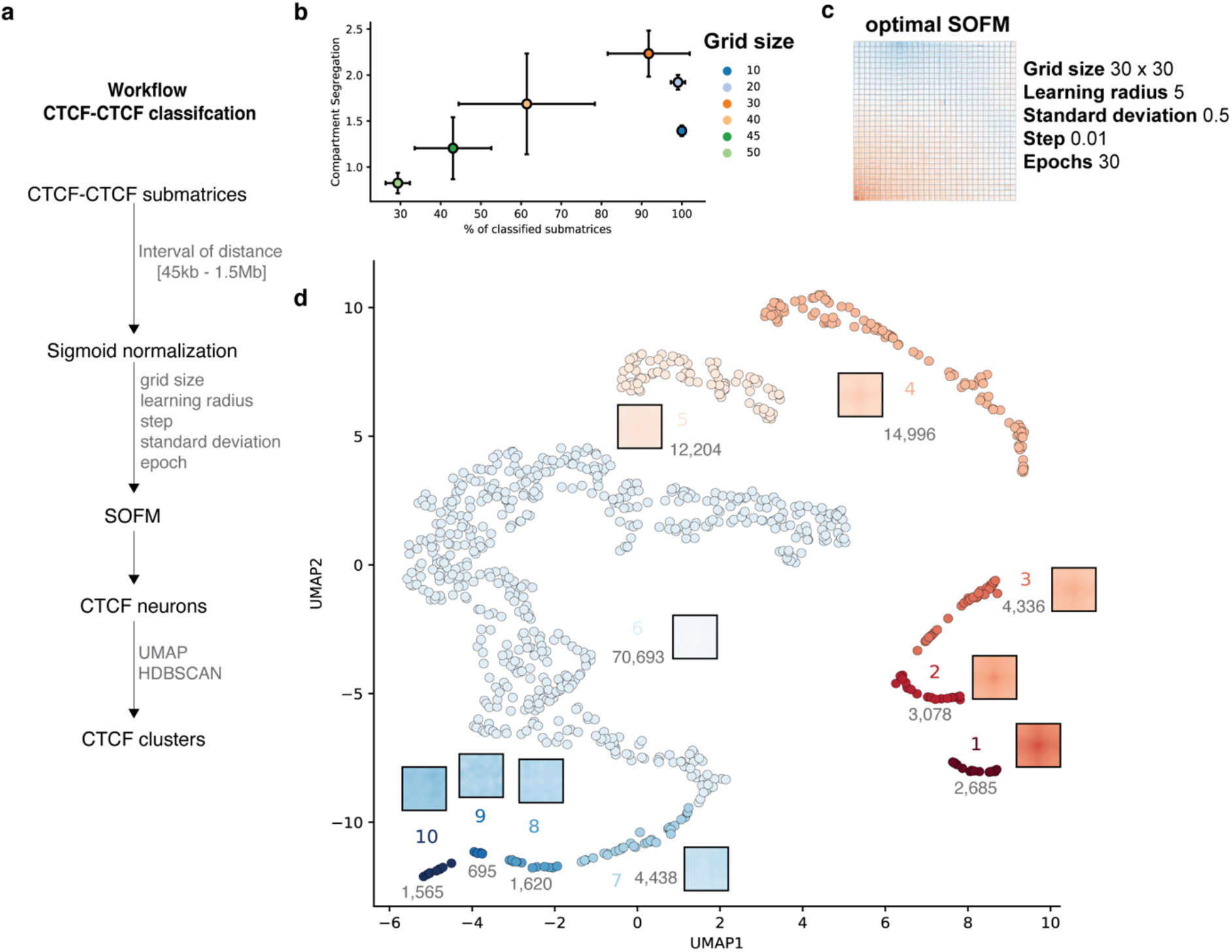
General overview of signal structure deconvolution. **a.** Schematic workflow to obtain CTCF clusters. **b.** SOFM parameters selected based on the percentage of classified matrices and the compartment segregation value (see also **Supplementary Fig. 1**). **c.** SOFM map showcasing the medoid of each neuron for optimal SOFM with the highest compartment segregation as well as highest percentage of classified CTCF-CTCF submatrices. **d.** Low dimensional representation of the SOFM neurons using a UMAP algorithm followed by a clustering method using HDBSCAN. A total of 10 clusters were obtained. Each point corresponds to a neuron in the SOFM map in panel c. Clusters are represented by the medoid signal and the number of CTCF-CTCF submatrices per cluster.

### Functional characterization of the CTCF-CTCF clusters

As described before, segregation of A/B compartment types between neurons was used as a metric to select the optimal set of parameters for the SOFM classifier. However, this measure was agnostic to the medoid submatrix topology used to cluster neurons. Interestingly, clusters with clear loop topology (clusters 1 to 5) are enriched in A compartments at the anchor points of the loops. Conversely, less structured clusters, with non-interacting CTCF-CTCF (clusters 6 to 10), are enriched in B compartments (**Figure 2a**). The separation between clusters 1 to 5 and 6 to 10 is also observed when revealing enrichment in CTCF pairs with different directionality (9), CTCF-CTCF clusters with clear interaction patterns are enriched in convergent-oriented CTCFs (clusters 1 to 4, **Figure 2b**), while non-interacting patterns are enriched in divergent-oriented CTCFs (clusters 7 to 10, **Figure 2c)**, and parallel-oriented CTCFs are slightly enriched in mid-interacting signal clusters (clusters 5 and 6, **Figure 2d**). Note that in these CTCF-CTCF orientation categories, significant enrichment in a category is usually accompanied by significant depletion in the others. Next, we observed that CTCF-CTCF clusters with clear loop topology (clusters 1 to 3) have a mean genomic distance between CTCF peaks of between 634 kb and 680 kb (**Figure 2e**), while clusters 7 to 10, more enriched in B compartment and divergent orientation, present shorter mean genomic distances spanning from ~510 kb to ~629 kb (**Figure 2e**). Note that this measure may be affected by our strict pre-selection of non-overlapping CTCF sites (**Materials and Methods**) resulting in a relatively sparse dataset, and, on average, more separated anchors.

**Figure 2.**
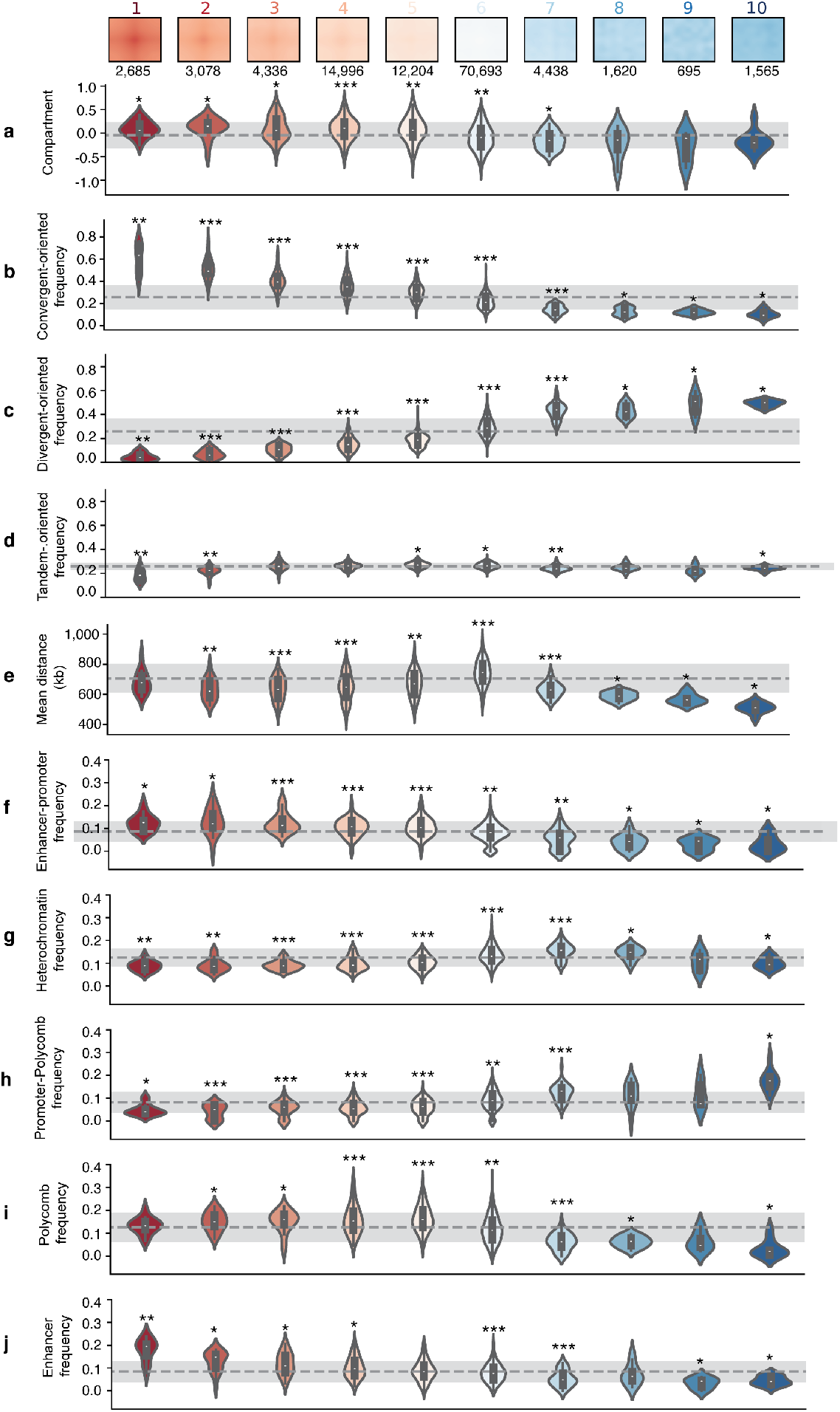
Distribution of multiple genomic features throughout CTCF clusters. In the first row the representative medoid of each CTCF cluster is represented. Below the different genomic features per cluster. In the first row the compartment type distribution, then the binding CTCF motif orientation, the distance between the two CTCFs and finally the chromatin state enrichment of the CTCF pairs. The dashed gray lines mark the mean, whereas the standard deviation is marked with lighter gray. Statistical significance against the mean is calculated using Wilcoxon test. p < 0.05 (*), p < 0.001 (**), p < 0.0001 (***).

To assess whether chromatin states correlate with the CTCF-CTCF clusters, we next measured chromatin state enrichment at CTCF sites for each cluster (**Materials and Methods**). Interestingly, pairs of CTCFs that form loop-like structures (clusters 1 to 4) are enriched with enhancer-promoter interactions (with one anchor point labeled as “enhancer” and the other as “promoter”, **Materials and Methods**) (**Figure 2f**). Anchors falling in the “heterochromatin” state have an almost opposite distribution: enriched in clusters 6 to 8 (**Figure 2g**). Interestingly, CTCF-CTCF cluster 10, which corresponds to the most B compartment cluster, present less heterochromatic anchors than expected but, in turn, is enriched in polycomb-promoter pairs (**Figure 2h**). CTCF-CTCF pairs with polycomb in both anchors are strongly enriched in mid-interacting clusters (clusters 4 to 6) and depleted in non-interacting pairs (clusters 7 to 10) (**Figure 2i**). Finally, enhancer-enhancer pairs, are more present in interacting clusters (clusters 1 to 4) and less abundant in non-interacting clusters (clusters 7 to 10) (**Figure 2j**). Interestingly cluster 1, with the strongest interaction pattern, presented the highest concentration of enhancer-enhancer loops, being this feature, it’s most distinct measure when compared to cluster 2 or 3.

Our analysis indicates the existence of clusters or types of CTCF-CTCF pairs that gradually expand from enhancer-promoter, convergent, mid-range interacting pairs in A compartment, to a polycomb-heterochromatin, divergent, short-range, non-interacting pairs in B compartment. Taken together, our results based solely in the interaction pattern of CTCF-CTCF proteins reveal two major types of CTCF-CTCF pairing that can further characterize well defined functional states.

### Loop calling using LOOPbit, a CNN trained with looping and non-looping CTCF-CTCF pairs

Next, we randomly subset a total of 6,000 CTCF-CTCF submatrices from clusters 1 to 5 as loop forming pairs (significantly enriched in promoter-enhancer loops) and another 6,000 CTCF-CTCF submatrices from clusters 7 to 10 as non-loop forming pairs. The two datasets, loop and non-loop, where next used to train a convolutional neural network (CNN). This CNN, that we called LOOPbit, aims at automatically assigning a probability of a CTCF-CTCF pair forming a loop in the central bin of a 9×9 submatrix extracted from genome-wide Hi-C experiments (**Figure 3a** and **Materials and Methods**). LOOPbit, which was trained by a 20% leave-out of the used data, was then assessed for accuracy and compared to other loop-calling methods using a recently published benchmark (23). LOOPbit, similarly to other published methods including HICCUPS (9), GOTHiC (35), HOMER (36), diffHic (37), HIPPIE (38), and Fit-Hi-C (39), detects an increasing number of chromatin loops in denser Hi-C experiments (**Figure 3b**). A solution to minimize this effect is to reduce data resolution (*i.e.*, from 5 kb to 40 kb) (**Supplementary Figure 2a**). We note here that LOOPbit is the loop caller with steeper slope, meaning that it is the most conservative in sparse dataset. Loops detected by LOOPbit were of similar average size as of HICCUPS, diffHiC and HIPPIE (~200 kb) and larger than those by GOTHiC (~80 kb) or HOMER (~100 kb) but shorter than those by Fit-Hi-C (~10 Mb) (**Figure 3c**). This loop size measure is again dependent on the resolution of the data as loops called at 40 kb resolution increased to ~1 Mb of size for all methods (**Supplementary Figure 2b**).

**Figure 3.**
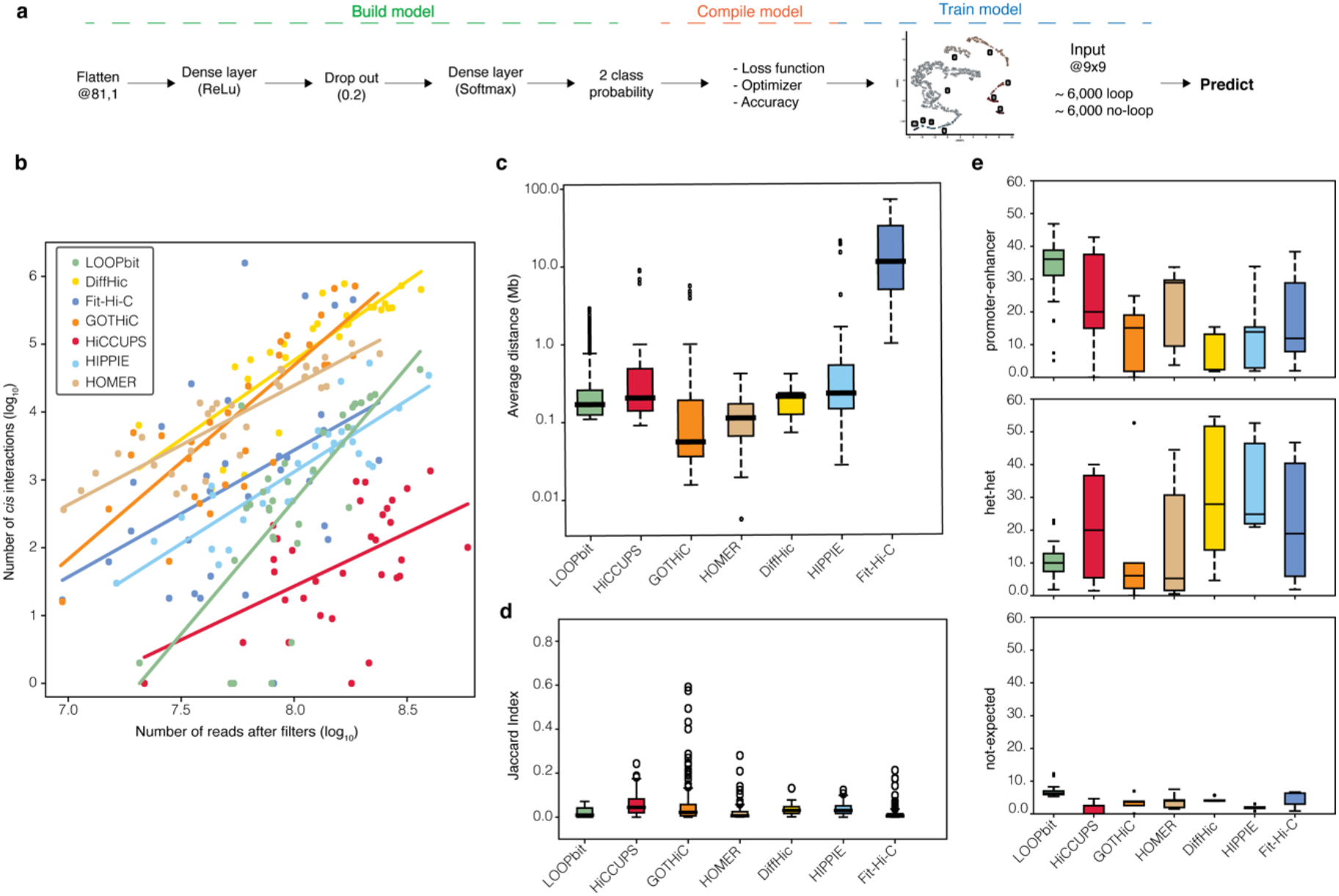
LOOPbit technical and biological benchmarking. **a.** CNN workflow, model building with the multiple layers to transform the input data. Model compilation to assess the accuracy and optimize the model. Finally, model training using the data coming from the CTCF-CTCF SOFM deconvolution. **b.** Representation of the number of reads after filters and the number of identified cis-interactions by LOOPbit in all experiments at a 5 kb resolution (n=32). **c.** Average distance between the identified loop-anchors of all the Hi-C experiments at 5 kb resolution (n=32). **d.** Boxplot representing the Jaccard Index, in here the overlapping between exact loop-anchors were considered to be the same loop between the same replicates (n=39). **e.** Proportion of the identified cis interactions based on the chromatin states at their anchoring points of the datasets at 5 kb resolution (n=32).

Next, we assessed the ability of LOOPbit to reproduce loop detection by measuring the Jaccard Index using replicates of previously published Hi-C experiments (**Materials and Methods** and **Supplementary Table 2**). Despite that LOOPbit yields continuous probability clouds instead of pinpointing single cells in the matrix, we used the exact same benchmark measures previously published (23) to calculate the Jaccard Index between replicates (that is to consider two loops identical if both their anchoring bins are the same). With such benchmark, LOOPbit results are slightly more reproducible in average than compared to HOMER and Fit-Hi-C, similar to GOTHiC, diffHiC and HIPPIE but lower than HICCUPS (**Figure 3d**). Finally, we compared the accuracy of the loop callers in terms of the biological relevance of the predicted loops. LOOPbit was able to detect higher percentage of enhancer-promoter loops than any other caller (**Figure 3e**, top panel). Concomitantly, LOOPbit also detects less loops between heterochromatic anchors (**Figure 3e**, middle panel) and has similar levels of non-expected loops (that is, between enhancer and heterochromatin, **Figure 3e**, bottom panel). Interestingly, and likely particular to LOOPbit as it was trained using CTCF-CTCF selected loops (**Materials and Methods**), ~56% of all detected loops in the training had an annotated CTCF site in both anchor points.

Altogether our results indicate that, although LOOPbit has a similar level of reproducibility as other loop callers, chromatin loops detected by LOOPbit show a clear enrichment in functional signatures when compared to the other methods.

## DISCUSSION AND CONCLUSION

In this work we introduce the use of the structure signal deconvolution in the context of colocalizing DNA-binding proteins. This methodology aims at identifying clusters of different structural patterns. Until now, most methods detecting structural patterns associated to DNA-binding proteins use aggregate peak analysis (APA) to show an average interaction pattern for different targets of interest. Unfortunately, APA is blind to small subsets with specific interaction patterns, subsets with different structures that could potentially be related to specific functions.

Here, we convoluted the genomic average CTCF-CTCF interaction pattern, and, based solely on this structural feature, were able to characterize distinct subpopulations. According to their structural pattern of interaction only, ten CTCF clusters were obtained (**Figure 1d**). Each CTCF-CTCF cluster has a specific genome compartment location and epigenetic state. The first observation is that the genomic distance between the CTCF pairs as well as their orientation are relevant features to classify CTCF-CTCF interactions between those that structurally form and do not form loops (1, 9). Also expected, but less evident is that loop forming clusters (that is, those from cluster 1 to cluster 5) are enriched in A compartment and drive principally enhancer-enhancer and enhancer-promoter interactions. In this category we noted that the first cluster, presenting the strongest pattern of interaction, and the largest genomic distances, was particularly enriched in enhancer-enhancer loops, suggesting transcriptional hubs with rosette like structures (40). On the other side of the spectrum, clusters of CTCF-CTCF pairs presenting sparse or blur signal of interactions are enriched in B-compartment type, heterochromatin or polycomb chromatin states and are spanning shorter genomic distance. These clusters are representative of either silent chromatin (41) or of polycomb-polycomb driven interactions (42, 43). Interestingly, we could capture the polycomb interacting network, which has been observed to be essential for cell differentiation and identity. Thus, cluster number 10, which is mostly associated to compartment B, no loop structure, few interactions, divergent CTCF binding sites and short genomic distances between anchor points, contained a surprising enrichment of promoter-polycomb states which could be explained by polycomb protecting a given promoter with CTCF from interacting with another CTCF site. While clusters 4 to 6, presented a loop pattern and were mostly associated to A compartment, were enriched in polycomb at both loop anchors, suggesting the formation of chromatin loops for a proper cell identity regulation. Together these findings define subcategories of CTCF-CTCF interactions within which only ~30% (clusters 1 to 5) are consistent with the most accepted model of CTCF loops bringing together promoters and enhancers to initiate transcription (14).

The deconvolution of the average signal between CTCF-CTCF pairs allowed us thus to identify two major groups, one forming a canonical loop (clusters 1 to 5) and a second segregating CTCF sites (clusters 7 to 10). This allowed us to generate a *bona fide* set of CTCF-CTCF submatrices that generate loops versus those that do not. Next, we used those two sets to train an CNN to develop the loop-caller LOOPbit, which was technically and biologically tested against several previously benchmarked methods (23). LOOPbit, which was applied to 33 Hi-C experiments at 5 kb resolution and 4 experiments at 40 kb resolution, resulted in a number and length of loops similar to all other benchmarked methods and suffered from the same limitations. Indeed, loop-callers cannot easily replicate findings when comparing replicates of low sequencing depths or binned at high resolutions. Fortunately, when high sequencing depth data is available, LOOPbit increases its reproducibility. Importantly, loops called by LOOPbit were found to be particularly relevant in terms of biological function. We found a clear enrichment of promoter-enhancer loops and depletion of loops between anchor points in heterochromatin state (9, 14). When compared to other callers, LOOPbit detected almost twice as many promoter-enhancer loops. This feature is likely to be a direct consequence of the SOFM classification and its ability to capture the functional information embedded in CTCF-pairings structures.

In summary, we have shown that signal deconvolution is able to define sub-classes of CTCF-driven chromatin loops with specific structural features. We then designed a software based on this classification and its loops were the most enriched in enhancer-promoter interactions among other loops called by other software. With this work we have characterized and validated the relationship between structure and function of CTCF driven chromatin loops and provide a method that can be trained to identify chromatin loops driven by other DNA binding proteins beyond CTCF.

## CODE AVAILABILITY

Meta-Waffle as well as LOOPbit are available on GitHub: (https://github.com/3DGenomes/metawaffle and https://github.com/3DGenomes/loopbit, respectively).

## FUNDING

This work was partially supported by the European Research Council under the 7^th^ Framework Program FP7/2007-2013 (ERC grant agreement 609989), the European Union’s Horizon 2020 research and innovation programme (grant agreement 676556) and the Spanish Ministerio de Ciencia e Innovación (BFU2017-85926-P). CRG acknowledges support from ‘Centro de Excelencia Severo Ochoa 2013-2017’, SEV-2012-0208 and the CERCA Programme/ Generalitat de Catalunya as well as support of the Spanish Ministry of Science and Innovation through the Instituto de Salud Carlos III and the EMBL partnership, the Generalitat de Catalunya through Departament de Salut and Departament d’Empresa i Coneixement, and the Co-financing with funds from the European Regional Development Fund (ERDF) by the Spanish Ministry of Science and Innovation corresponding to the Programa Opertaivo FEDER Plurirregional de España (POPE) 2014-2020 and by the Secretaria d’Universitats i Recerca, Departament d’Empresa i Coneixement of the Generalitat de Catalunya corresponding to the programa Operatiu FEDER Catalunya 2014-2020. Funding for Open Access Spanish Ministerio de Ciencia, Innovación y Universidades (BFU2017-85926-P).

## ACKNOWLEDGMENTS

We thank the authors of (23) for providing the dataset used to make possible the comparison of LOOPbit with other loop callers. We thank David Castillo, Luciano Di Croce, Sandra Peiró and François Le Dily for their continuous discussions and support to the development of Meta-Waffle and LOOPbit.

## CONFLICT OF INTEREST

None declared.

## Notes

### Competing Interest Statement

The authors have declared no competing interest.

