## Supplementary Material for "Identification of chromatin loops from Hi-C interaction matrices by CTCF-CTCF topology classification"

**Supplementary Table 1.** GM12878 Hi-C experiments used to deconvolve the CTCF-CTCF topology (9).

| ID | Reestriction enzyme | Filtered reads |
|---|---|---:|
| GSM1551552 | MboI | 361,207,349 |
| GSM1551553 | MboI | 136,507,541 |
| GSM1551554 | MboI | 255,191,180 |
| GSM1551555 | MboI | 129,354491 |
| GSM1551556 | MboI | 148,151,521 |
| GSM1551557 | MboI | 172,219,884 |
| GSM1551558 | MboI | 99,044,057 |
| GSM1551559 | MboI | 48,503,486 |
| GSM1551560 | MboI | 48,052,662 |
| GSM1551561 | MboI | 115,277,508 |
| GSM1551562 | MboI | 58,351,032 |
| GSM1551563 | MboI | 209,244,601 |
| GSM1551564 | MboI | 94,795,755 |
| GSM1551565 | MboI | 94,272,224 |
| GSM1551566 | MboI | 147,025,950 |
| GSM1551567 | MboI | 119,160,683 |
| GSM1551571 | MboI | 228,444,864 |
| GSM1551572 | MboI | 218,414,758 |
| GSM1551573 | MboI | 77,453,007 |
| GSM1551577 | MboI | 54,523,851 |
| GSM1551578 | MboI | 121,751,924 |
| GSM1551588 | DpnII | 58,470,721 |
| GSM1551589 | DpnII | 76,770,682 |
| GSM1551590 | DpnII | 61,529,714 |
| GSM1551591 | DpnII | 93,923,134 |
| GSM1551598 | MboI | 82,406,062 |

**Supplementary Table 2.** Hi-C experiments used for LOOPbit benchmarking. All the experiments were pre-processed and filtered using TADbit (28) and OneD normalized (29).

| Dataset | ID | Restriction enzyme | Cell type | Filtered reads | Resolution (kb) |
|---|---|---|---|---:|---|
| JIN | GSM1055805 | HindIII | H1-hESC | 134,678,276 | 5 |
| JIN | GSM1055800 | HindIII | IMR90 | 104,929,328 | 5 |
| JIN | GSM1055801 | HindIII | IMR90 | 175,071,658 | 5 |
| JIN | GSM1154021 | HindIII | IMR90 | 97,395,328 | 5 |
| JIN | GSM1154022 | HindIII | IMR90 | 79,559,232 | 5 |
| JIN | GSM1154023 | HindIII | IMR90 | 52,425,794 | 5 |
| JIN | GSM1154024 | HindIII | IMR90 | 54,082,516 | 5 |
| RAO | GSM1551552 | MboI | GM12878 | 361,207,349 | 5 |
| RAO | GSM1551569 | MboI | GM12878 | 72,934,660 | 5 |
| RAO | GSM1551570 | MboI | GM12878 | 77,651,974 | 5 |
| RAO | GSM1551571 | MboI | GM12878 | 228,444,864 | 5 |
| RAO | GSM1551572 | MboI | GM12878 | 218,414,758 | 5 |
| RAO | GSM1551573 | MboI | GM12878 | 77,453,007 | 5 |
| RAO | GSM1551574 | MboI | GM12878 | 81,318,602 | 5 |
| RAO | GSM1551575 | MboI | GM12878 | 80,613,339 | 5 |
| RAO | GSM1551576 | MboI | GM12878 | 80,438,152 | 5 |
| RAO | GSM1551577 | MboI | GM12878 | 54,523,851 | 5 |
| RAO | GSM1551578 | MboI | GM12878 | 121,751,924 | 5 |
| RAO | GSM1551587 | DpnII | GM12878 | 63,854,975 | 5 |
| RAO | GSM1551588 | DpnII | GM12878 | 58,470,721 | 5 |
| RAO | GSM1551589 | DpnII | GM12878 | 76,770,682 | 5 |
| RAO | GSM1551590 | DpnII | GM12878 | 61,529,714 | 5 |
| RAO | GSM1551591 | DpnII | GM12878 | 93,923,134 | 5 |
| rao | GSM1551599 | MboI | IMR90 | 164,365,813 | 5 |
| rao | GSM1551600 | MboI | IMR90 | 181,640,359 | 5 |
| rao | GSM1551601 | MboI | IMR90 | 20,676,015 | 5 |
| rao | GSM1551602 | MboI | IMR90 | 90,970,187 | 5 |
| rao | GSM1551603 | MboI | IMR90 | 186,707,523 | 5 |
| rao | GSM1551604 | MboI | IMR90 | 198,492,270 | 5 |
| rao | GSM1551605 | MboI | IMR90 | 216,948,852 | 5 |
| Dixon 2015 | GSM1267196 | HindIII | H1-hESC | 172,971,685 | 5 |
| dixon 2015 | GSM1267197 | HindIII | H1-hESC | 103,476,074 | 5 |
| Sexton | GSM849422 | DpnII | Fly embryo | 31,357,023 | 40 |
| dixon 2012 | GSM862723 | HindIII | H1-hESC | 21,292,727 | 40 |
| dixon 2012 | GSM892306 | HindIII | H1-hESC | 134,678,276 | 40 |
| dixon 2012 | GSM862724 | HindIII | IMR90 | 102,906,483 | 40 |
| dixon 2012 | GSM892307 | HindIII | IMR90 | 104,974,904 | 40 |

**Supplementary Table 3.** CTCF ChIP-seq experiments used in the analysis.

| Experiment | Cell type | Accession number |
|---|---|---|
| ctcf cHip-SEQ | H1-hESC, GM12878 | GSE29611 |
| ctcf cHip-SEQ | IMR90 | GSE31477 |
| ctcf cHip-SEQ | Embryo 14-16hr Oregon-R | GSE47264 |

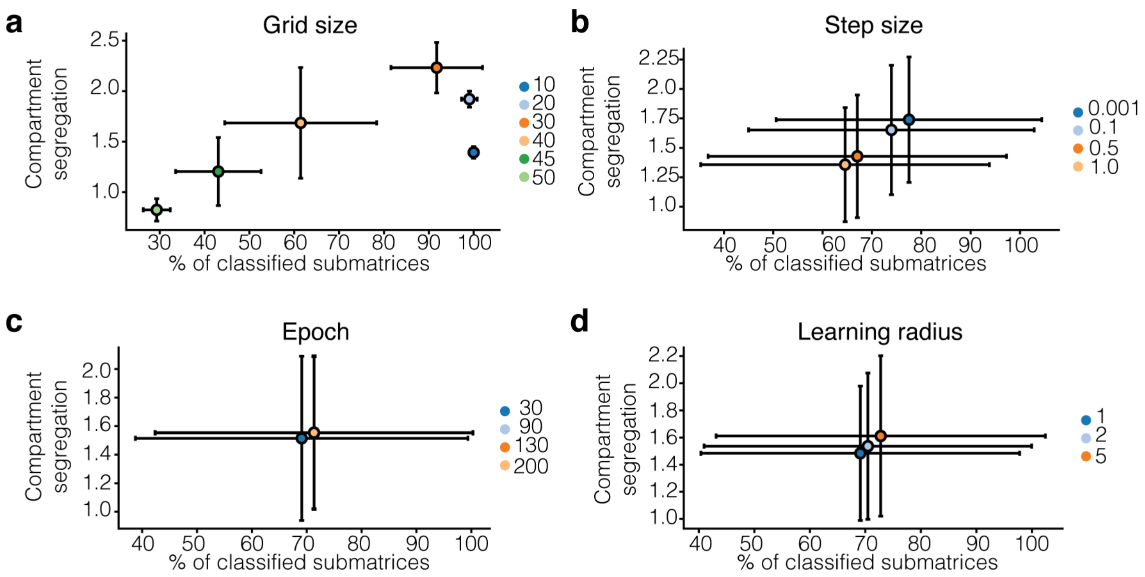


**Supplementary Figure 1. SOFM parameters.** SOFM parameters, grid size, step size, epoch and learning radius, based on the percentage of classified matrices and the compartment segregation value (see also **Fig. 1**), **a**, **b**, **c** and **d,** respectively.

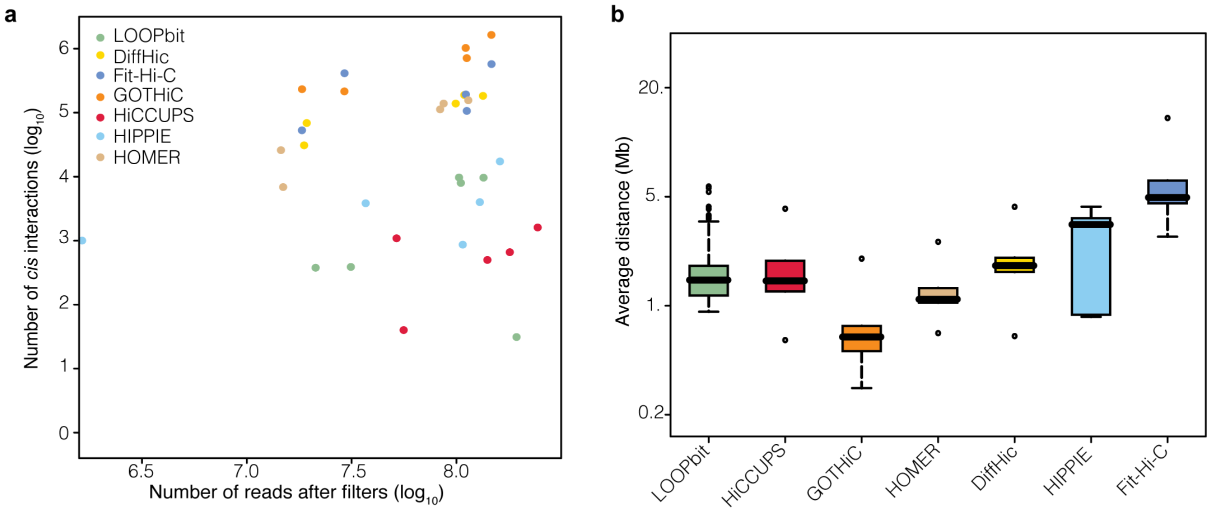


**Supplementary Figure 2. Results benchmark at 40 kb resolution. a.** Number of reads after filters and the number of identified cis-interactions by LOOPbit in all experiments at a 40 kb resolution (n=5). **b.** Average distance between the identified loop-anchors of all the Hi-C experiments at 40 kb resolution (n=5).

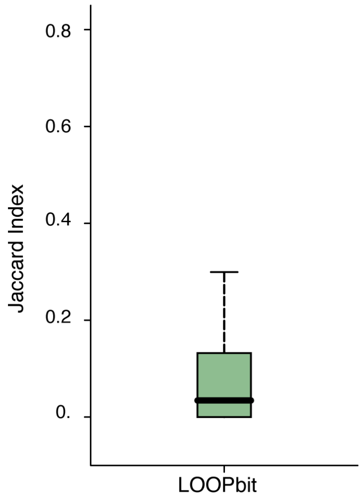


**Supplementary Figure 3. Jaccard Index allowing 70% of overlapping between chromatin loops.** Boxplot representing the Jaccard Index, in here an overlap of 70% between chromatin loops were considered to be the same loop between the same replicates (n=39).
